# Altered functional connectivity and spatiotemporal dynamics in individuals with sleep disorders

**DOI:** 10.1101/2024.08.28.610147

**Authors:** Lauren Daley, Prabhjyot Saini, Harrison Watters, Yasmine Bassil, Eric H. Schumacher, Lynn Marie Trotti, Shella Keilholz

**Author notes:** these authors contributed equally to this work (Lynn Marie Trotti, Shella Keilholz).

## Abstract

Idiopathic hypersomnia (IH) is a sleep disorder characterized by highly disruptive symptoms. Like narcolepsy type 1, a well-characterized sleep disorder, individuals with IH suffer from excessive daytime sleepiness, though there is little overlap in metabolic or neural biomarkers across these two disorders. This lack of common pathophysiology, combined with the clear overlap in symptoms presents an ideal paradigm for better understanding the impact of IH on an individual’s functional activity and organization, and potentially, the underlying pathophysiology. This study examines the observed functional connectivity in patients with IH, and patients with narcolepsy type 1 (NT1) against healthy control individuals. Static functional connectivity is compared, as are quasi-periodic patterns, acquired from the BOLD timecourse, for all groups. In addition to baseline data comparison, the study also included a post-nap condition, where the individuals included in this analysis napped for at least 10 minutes prior to the scanning session, to explore why individuals with IH do not feel “refreshed” after a nap like individuals with NT1 do. Assessing the groups’ spatiotemporal patterns revealed key differences across both disorders and conditions: static connectivity revealed at baseline higher subcortical connectivity in the NT1 group. There was also observably less connectivity in the IH group both at baseline and post-nap, though none of these static analyses survived multiple comparisons correction to reach significance. The QPP results however found significant differences in the IH group in key networks, particularly the DAN/FPCN correlation is significantly different at baseline vs. post-nap, a trend not observed in either the control or NT1 groups. The DAN and FPCN are drastically altered both at baseline and post-nap when compared to the other groups, and may likely be a disorder-specific result. This study demonstrates that key networks for arousal are more heavily disrupted in IH patients, who are less affected by a nap, confirmed through both subject reporting and functional evidence through spatiotemporal patterns.

## Introduction

Sleep disorders are highly disruptive and detrimental to an individual’s health and well-being, mitigated only through accurate diagnosis and effective treatment (Ramar et al. 2021). However, accurate diagnosis can be challenging due to overlapping symptoms across disorders. Excessive daytime sleepiness (EDS) is a primary feature in sleep disorders (e.g. IH, NT1/NT2), and secondary feature in psychiatric (e.g. depression) neurologic (e.g. Parkinsons’s Disease), and medical disorders (e.g hepatic encephalopathy). While the pathophysiology of narcolepsy type 1 (NT1) is well characterized by the loss of hypocretin neurons, idiopathic hypersomnia (IH) remains unclear, with few evidence-based treatment options (Maski et al. 2021). IH is characterized by EDS despite normal or long nighttime sleep duration, often accompanied by long, unrefreshing naps and difficulty awakening (Trotti 2017; Miglis et al. 2020).

Both IH and NT1 are characterized by EDS despite sufficient duration of nighttime sleep. However, they differ with respect to sleep quality (poorer in people with NT1) (Takahashi 2003) and the quality of daytime wakefulness (Ramm et al. 2019; Dauvilliers et al. 2019). People with NT1 tend to experience sudden attacks of sleepiness, whereas people with IH tend to have more persistent, less irresistible daytime sleepiness (Miglis et al. 2020). Another major difference in the two disorders is the impact of daytime naps. People with NT1 tend to feel refreshed from a short nap, while people with IH tend to take long, unrefreshing naps that result in a temporary worsening of symptoms, known as severe sleep inertia (Arnulf, Leu-Semenescu, and Dodet 2022). Similarly, people with IH have a much more difficult time awakening from night sleep (often referred to as “sleep drunkenness”) (Rassu et al. 2022). In addition, unlike NT1, for which cerebrospinal fluid hypocretin (orexin) deficiency is a reliable biomarker reflecting disease pathology (Huang et al. 2018), no definitive biomarkers have been definitively established for IH, though several mechanisms have been proposed such as a positive allosteric modulator at the GABAA receptor (Trotti et al. 2015; Lu, Jhou, and Saper 2006). Lack of a biomarker renders an accurate and timely diagnosis difficult to achieve (Miglis et al. 2020). Similarly, little is known about how brain activity is altered in individuals with IH, though individuals with NT1 are known to have altered neural connectivity (Wu et al. 2022).

Functional magnetic resonance imaging (fMRI), is a tool commonly used to characterize, diagnose, and assess neurological disorders. When a patient group is compared against a healthy baseline using fMRI, the acquired images produce significant spatial differences in whole-brain connectivity and activation levels. The variation from baseline can either aid in understanding/characterizing the patient group’s affliction (such as movement disorders including Parkinson’s disease and Huntington’s disease, or cognitive conditions including Alzheimer’s disease), or act as a diagnostic tool in the case of some often outwardly obscured disorders, especially when fMRI is performed in conjunction with other neuroimaging modalities (Specht 2020; Salvador et al. 2019). Because the cause or causes of IH are currently unknown, analyzing resting state fMRI data from people with IH may help to elucidate the disorder’s neural underpinnings and how its pathophysiology translates to neural dynamic activity.

The current diagnosis regimen for both NT1 and IH is a clinical test called the Multiple Sleep Latency Test (MSLT), designed to measure how quickly an individual can fall asleep in a controlled daytime environment, and is often recommended by a physician if a patient reports EDS. But very little is revealed through this test about the *cause* of the patient’s sleepiness, so pairing these diagnoses with functional imaging may differentiate the origin of sleepiness between a patient with NT1 and a patient with IH (Arand and Bonnet 2019).

Existing functional neuroimaging studies of IH and NT1, although limited, have found evidence of altered global neural network connectivity, specifically in default mode network (DMN) and task positive network (TPN), compared to non-sleepy control subjects. The DMN in particular was found to be significantly lower in activation in people with sleep disorders (both IH and NT1) (Pomares et al. 2019; Fulong et al. 2020). This network is typically less active during tasks and goal-oriented situations, while remaining active at rest (Raichle et al. 2001; van Es et al. 2023). The DMN is typically linked to higher-order cognitive processes like internally focused attention or mind wandering (Bola and Borchardt 2016; Raichle et al. 2001; Pomares et al. 2019). High anticorrelation between the DMN and TPN is a key characteristic of healthy, “normal” functional connectivity, and significantly altered DMN activation is often seen in individuals with neurological disorders, including those related to sleep (Raichle et al. 2001; Järvelä et al. 2020). A consistently active DMN is often considered to indicate alertness or arousal; activity in the DMN decreases during sleep (Satpute and Lindquist 2019). Conversely, the TPN is typically more active during tasks and goal-oriented thinking/actions and has key nodes in both the frontoparietal network (FPN) and dorsal attention network (DAN) (Di and Biswal 2014). DMN/TPN anticorrelation is thought to be an inherent property of neural network-level activity and can be used to measure arousal level in fMRI participants (Di and Biswal 2014; Satpute and Lindquist 2019).

To date, IH studies have only measured time-averaged functional connectivity, which is not sensitive to the variation of activity over time that is captured by newer time-resolved analysis methods. Preserving this temporal data further elucidates resting-state dynamic neural activity (Bolt et al. 2022; Keilholz 2023; Majeed et al. 2011; Liu et al. 2018; Kiviniemi et al. 2011; Allen et al. 2014) and may prove more sensitive to alterations that occur in brain disorders. In fact, one study from Du et al conducted a sliding-windowed analysis on fMRI data acquired from individuals diagnosed with schizophrenia, and identified some targeted DMN regions that are significantly impaired as a result of the disorder – a result obscured in the static results (Du et al. 2016). One dynamic method, quasi-periodic pattern (QPP) detection, has been successfully used to identify widespread and localized pattern variation across pathologies (Abbas, Bassil, and Keilholz 2019; Belloy, Shah, et al. 2018). QPPs are repeating spatiotemporal patterns in neural activity in DMN and TPN (Majeed et al. 2011) and have been observed in the BOLD signal, correlated to infraslow neural activity (Thompson et al. 2014; Grooms et al. 2017).

QPPs are ideal for a network-level analysis over time as they preserve temporal information, do not distort or change the original data, and can be used to quantify whole-brain spatiotemporal activity. The DMN and TPN are known to be involved in arousal (Satpute and Lindquist 2019; Di and Biswal 2014), and appear more anticorrelated to each other in high arousal state(s). In low arousal state(s), complementary datasets have demonstrated lesser anticorrelation, while maintaining an equally active DMN, and a dampened TPN level of activity (Wang et al. 2016; Abbas et al. 2019; Belloy, Naeyaert, et al. 2018). Differences in QPPs have been shown in other neurological disorders, including ADHD, for which Abbas et al showed evidence that individuals with ADHD have disrupted DMN/TPN connectivity as compared to a control (Abbas, Bassil, and Keilholz 2019). They also found that individuals with ADHD have an overall weaker QPP, as it contributes less to their overall functional connectivity, which are findings consistent with static FC studies in subjects with ADHD (Castellanos et al. 2008; Uddin et al. 2008).

This study aims to investigate functional connectivity of individuals with IH, or NT1 vs. non-sleepy control participants to determine whether differences in QPPs between IH patients and non-sleepy controls reflect disease-independent effects of EDS or disease-specific patterns of dysfunction that might identify disease mechanisms in IH or lead to the future development of more accurate diagnostic testing. We hypothesize that, although both IH and NT1 patients have pathologic daytime sleepiness, their QPPs (amplitude and inter-network correlationi values) will differ, reflecting the different pathophysiology and symptom expression of these two disorders. We further hypothesize that analyzing QPPs will isolate the different effects of a mid-length nap in these groups. To investigate these two central hypotheses, we analyzed rs-fMRI data from three groups (IH, NT1, control) before and after a nap using QPPs, focusing on the DMN and TPN regions and the network correlations. These results were validated with sensitivity analyses, global signal power spectra, and subject controls.

## Methods

### Participants selection

Participants with IH and NT1 were recruited from the Emory Sleep Center and from patient advocacy/support groups. Non-sleepy controls were recruited from advertisements around the Emory campus and advocacy/support groups. To be eligible for participation, IH and NT1 participants had to meet International Classification of Sleep Disorders, 3^rd^ edition (Sateia 2014), diagnostic criteria for their disorder and were asked to discontinue any IH or NT1 treatment for at least five half-lives prior to imaging. Control participants had to report habitual sleep durations of 6.5 to 9.0 hours, deny problematic sleepiness and have normal scores (≤10) on the Epworth Sleepiness Scale, have overnight polysomnography without nocturnal sleep pathology (i.e., obstructive sleep apnea with apnea hypopnea index > 10 or frequent periodic limb movements with arousals), and have a multiple sleep latency test with a normal mean sleep latency of 8 minutes or higher. Patient and control participants also were free of serious or unstable psychiatric disorders and major neurologic disorders. All participants provided written informed consent and this protocol was approved by the Emory University Institutional Review Board.

### MRI Acquisition

The dataset was acquired at the Biomedical Imaging Technology Center (Emory University) on a 3T Siemens MR scanner. An anatomic reference scan was acquired for each patient, with a T1-weighted MPRAGE sequence, with TR = 2300 ms, TE = 2.75 ms, slice thickness=0.8mm, and flip angle=8°. Resting-state fMRI scans were acquired using an interleaved EPI sequence, with TR=2s, TE=28ms, slice thickness=3.3mm, and flip angle=90°. Each subject was scanned on two separate days: one day with no nap prior to acquisition, and then one day immediately after a nap of a duration of at least 10 minutes. The day with no nap served as a waking baseline, while the day with a nap was an assessment of potential effects of napping. Participants were counterbalanced for the order of nap/no-nap days.

### EEG Acquisition

EEG was collected via BrainVision 64 channel EEG cap using BrainVision Analyzer software (“BrainVision Analyzer | Brain Products GmbH > Solutions” 2020), applied prior to both MRI acquisition sessions. On nap-day, the nap opportunity occurred in a dark, quiet room outside the scanner and was monitored with EEG recordings. Naps were allowed to continue up to approximately 30 minutes of sleep duration, within the limitations of the MRI schedule and each participant’s sleep onset latency. Participants were awoken from the nap and immediately transported via wheelchair into the MRI scanner (to minimize effects of physical exertion on the rate of dissipation of sleep inertia). Once in the scanner, participants first underwent a 10-minute working memory scan (results reported separately) and then the 10-minute resting state scan described above. Subjects who were unable to nap for at least 10 minutes were removed from the conditional comparisons, as well as subjects who were only scanned on one day. Participants were also instructed to not fall asleep in the scanner while being scanned. EEG monitoring was continued throughout the scan, but because of technical limitations, simultaneous EEG-MRI data were only available on a subset of participants

### Preprocessing and QPP Detection

The fMRI and anatomical scans were preprocessed with CPAC pipeline (Craddock et al. 2013). The anatomical scans were preprocessed using N4 bias correction, skull-stripping, and linear and non-linear registration. Functional scans were preprocessed with slice-time correction, distortion correction, motion correction, nuisance signal regression, global signal regression, temporal filtering (BW=0.01-0.1 Hz) and registration. Participants whose imaging scans did not meet preprocessing quality requirements (e.g., excessive head motion or signal dropout) were excluded from the analyses (n = 1 with narcolepsy, n = 2 non-sleepy control). The scans were registered using the Brainnetome atlas, a human brain atlas with structural and functional parcellations (210 cortical regions, 36 sub-cortical regions) (Fan et al. 2016). All image voxels were sorted into these regions and subsequently Yeo’s networks. To validate previous studies that only performed time-averaged connectivity analyses, conventional functional connectivity was also visualized, using Pearson correlation coefficients. A t-test (p-value>0.05) was used to determine significance for these values, corrected for multiple comparisons with a multiple comparisons correction using false discovery rate (Lee and Lee 2018; “False Discovery Rate” 2016).

QPPs were acquired from the preprocessed functional time series using an adapted version of the pattern-finding algorithm developed by *Majeed et al.* for a window length of 20 seconds (Majeed et al. 2011; Abbas et al. 2019), schematized in Figure 1. This algorithm uses time-series data acquired during BOLD-fMRI for an entire group (all subjects concatenated) to produce an initial template of a repeating spatiotemporal pattern. The algorithm then iterates through every single datapoint to update the template, until convergence. The output QPP template most accurately represents underlying infraslow activity dynamics (Majeed et al. 2011; Thompson et al. 2014; Abbas et al. 2019). This template, thought to be representative of the entire group, is overlaid onto the entire averaged time-course to measure how each time-point correlates with it, visualized both in single-cycle waveform plots and heatmap visualizations.

**Figure 1:**
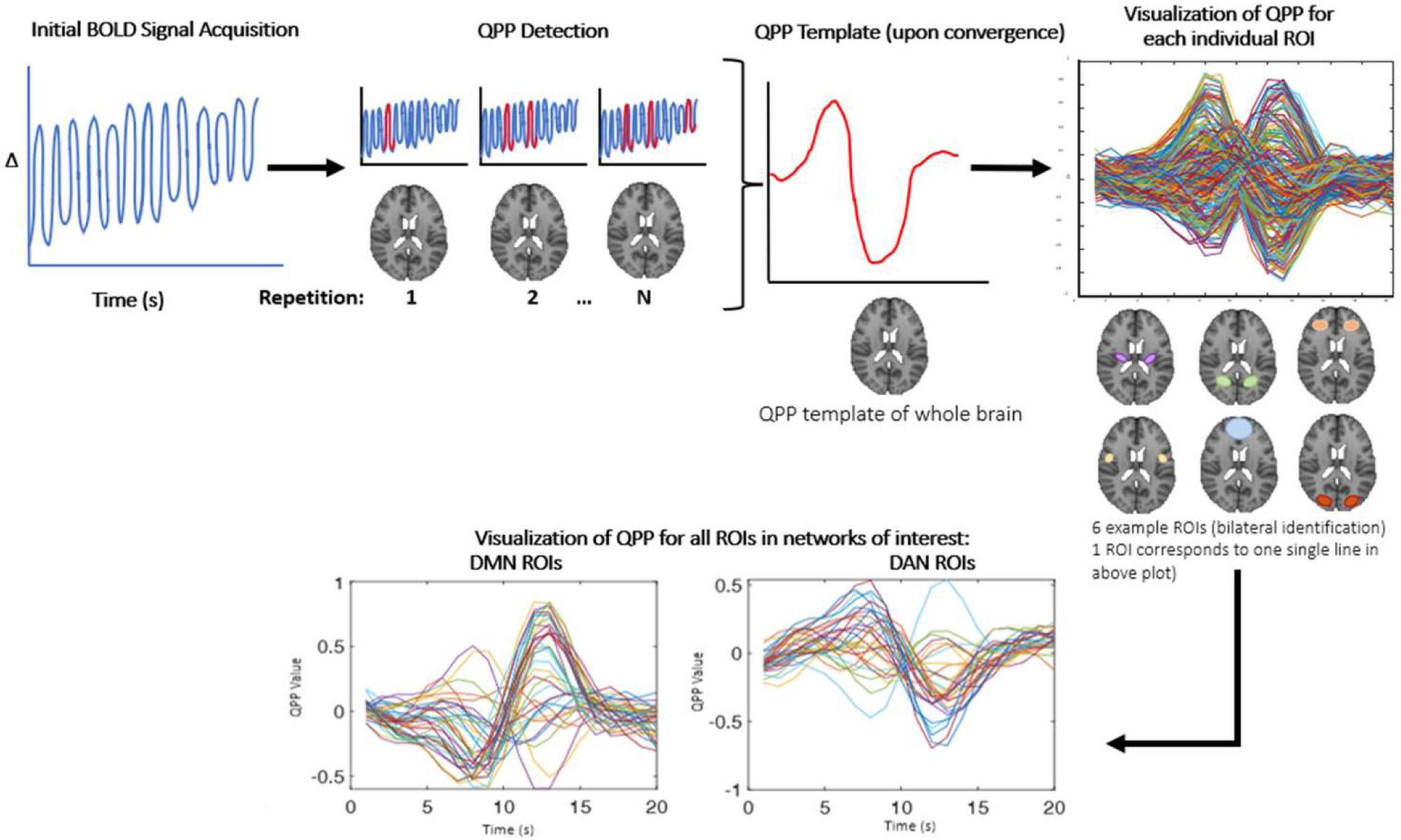
QPPs were detected in a method adapted from Majeed et al, as described in the above figure. The initial BOLD signal (far left plot) is input to the QPP detection algorithm, which then chooses a random starting (time)point, and develops an initial template of the QPP (indicated by the red sections in the QPP detection panel). This template is then compared against every single timepoint for correlation, and if it surpasses the threshold, the template is then updated to reflect this. This is iteratively done until convergence, and the template is the final QPP (shown above in red). To visualize the QPP, this study used the Brainnetomme atlas, and therefore produced 246 individual QPPs for each ROI – for sake of clarity, only 6 sample ROIs are shown in the above figure’s brain atlas. Based on atlas labels, these ROIs are then divided into network-specific patterns, later averaged to form one single line, as shown in later results (for example, figure 7).

### QPP Analyses

QPP template voxel-wise Pearson correlation values were calculated and plotted for each condition, followed by heatmap visualizations of the activity in each parcel over time in the template (see figure 3, panel a). QPP regression was also performed, and results were compared to non-regressed functional connectivity, to assess the pattern’s contribution to overall activation. Waveform depictions of a single-cycle QPP were produced for each unique condition (see figure 6). For network-level waveform visualizations, networks of interest include the DMN (denoted as ‘default mode’), dorsal attention, ventral attention, and fronto-parietal networks. Both the dorsal attention and fronto-parietal are subnetworks of the TPN, and as such we expect to see some level of anticorrelation between them and the DMN. These were used to assess how the detected QPP differs spatially across nap and no-nap conditions intra-group and inter-group. Correlation values between networks of interest were calculated with Pearson’s correlation coefficients and then tested for significance using a Fisher’s confidence test (Diedenhofen and Musch 2015). To more quantitatively examine how QPPs differ across conditions, the squared difference of the DMN/TPN correlation was calculated for each group: the difference in DMN/TPN correlation values at baseline and post-nap was calculated, then squared, then plotted on the same axes.

For each group, the QPP was acquired twice: once by concatenating all individuals’ data within the algorithm, thus retrieving the “group” QPP; the second time by inputting only one individual’s data into the algorithm at a time, thus retrieving the “individual” QPP, which was then averaged across all subjects post-QPP detection. Correlation values for key networks are included in the results table 1 for both QPP values. We also examined the inter-network correlation individual QPP using a Kruskal-wallis test for multiple comparisons. For the individual QPPs’ inter-network correlation values, p-values were calculated using the Kruskal-wallis test for each comparison (three-way comparison of all groups at baseline, three-way comparison of all groups post-nap). The DMN/TPN correlation value was compared across all three groups to determine any resultant significance both at baseline and post-nap.

**Table 1:** Includes correlation values between the networks of interest (default mode network (DMN) with dorsal attention network (DAN), frontoparietal network (FPCN), task-positive network (TPN), DAN with FPCN) at baseline for all groups; both group QPP and individual (italicized) QPP average and standard deviation (italicized). Of note, the group QPP and individual QPP averages are highly varied, with little agreement within groups or conditions. To denote significance, asterisks are used: *significance within group across conditions, **significance across groups within 1 condition.

| Group | Condition | Correlation Value (AU) |  |  |  |
| --- | --- | --- | --- | --- | --- |
|  |  | DMN/DAN | DMN/FPCN | DMN/TPN | DAN/FPCN |
| control | baseline | -0.87<br>*-0.57±0.56* | 0.98<br>*0.44±0.49* | 0.20<br>*-0.34±0.57* | -0.80<br>*-0.15±0.63\*\** |
|  | post-nap | -0.93<br>*-0.64±0.22* | 0.97<br>*0.20±0.38* | 0.28<br>*-0.39±0.37\*\** | -0.91<br>*<u>-0.14±0.45</u>* |
| IH | baseline | -0.70<br>*-0.35±0.49* | 0.9476<br>*<u>0.58±0.29</u>* | 0.33<br>*0.02±0.44* | -0.46*<br>*0.049±0.29\*\** |
|  | post-nap | -0.92<br>*-0.34±0.53* | 0.98<br>*0.36±0.44* | -0.41<br>*-0.064±0.48\*\** | -0.89*<br>*0.35±0.44* |
| NT1 | baseline | -0.26<br>*-0.067±0.53* | 0.55<br>*0.46±0.31* | -0.031<br>*0.17±0.48* | 0.59<br>*0.52±0.35\*\** |
|  | post-nap | -0.53<br>*-0.10±0.50* | 0.83<br>*0.63±0.34* | 0.016<br>*0.24±0.49\*\** | -0.029<br>*0.17±0.38* |

### Global Signal Validation

Global signal has previously been linked to individual variance in QPPs and levels of arousal, determined by EEG (Yousefi et al., 2018; Wong et al., 2013). To develop a better understanding of how each group’s neural dynamics differ and change on an individual and group basis, we conducted a simple global signal validation check, based on the method used by Anumba et al. (Anumba et al. 2023). A z-scored mean of global signal was calculated, followed by power spectra for each group with a 95% confidence interval. Both peak amplitude and overall trend were analyzed.

### Sensitivity Check

Because some of the subjects are affected by sleep disorders, there was a strong possibility of them involuntarily falling asleep in the scanner, which would no longer be categorized as resting-state fMRI data. Thus, to evaluate for this possible confound, we repeated our analyses limited to those whose EEG demonstrated sustained wakefulness throughout the fMRI scan, when sufficient data were available.

## Results

### Participants

Participants had diagnoses of IH (n = 15) or NT1 (n = 11) or were non-sleepy controls (n = 10). They had a mean age of 33.3 years (+/− standard deviation 10.2) and twenty-five (69.4%) were women; neither age nor gender differed by diagnosis. As anticipated, those with IH or NT1 were significantly sleepier than controls, by Epworth sleepiness scale (15.8 +/− 3.3 for IH, 17.6 +/− 3.6 for NT1, 6.2 +/− 2.9 for controls, p <0.0001, IH=NT1>controls) and multiple sleep latency test mean latency (5.3 +/− 1.7 for IH, 2.2 +/− 1.4 for NT1, 13.4 +/− 3.1 for controls, p < 0.0001, NT1<IH<controls).

### Baseline Comparison

We first performed analyses on only the baseline condition data when subjects did not have a nap prior to scanning. For the baseline condition, all 36 participants were included in analyses. Time-averaged functional connectivity analyses were performed first, and half connectivity matrices can be seen in Figure 2(a), for Yeo’s 7 networks and subcortical regions. Overall, the control group visually demonstrates more mean functional connectivity compared to the IH and NT1 groups. The NT1 group showed more differences in the subcortical areas compared to the other groups. Some networks demonstrate minor group-level differences. To test for statistical significance, a network-level t-test (p<0.05) was performed between all paired groups’ (IH-control, IH-NT1, NT1-control) connectivity differences; these results are in figure 2(b). After controlling for multiple comparisons, no group pairing demonstrated any significant difference, though there were the most non-controlled differences observed in the IH-control pairing at both the ROI and network level.

**Figure 2:**
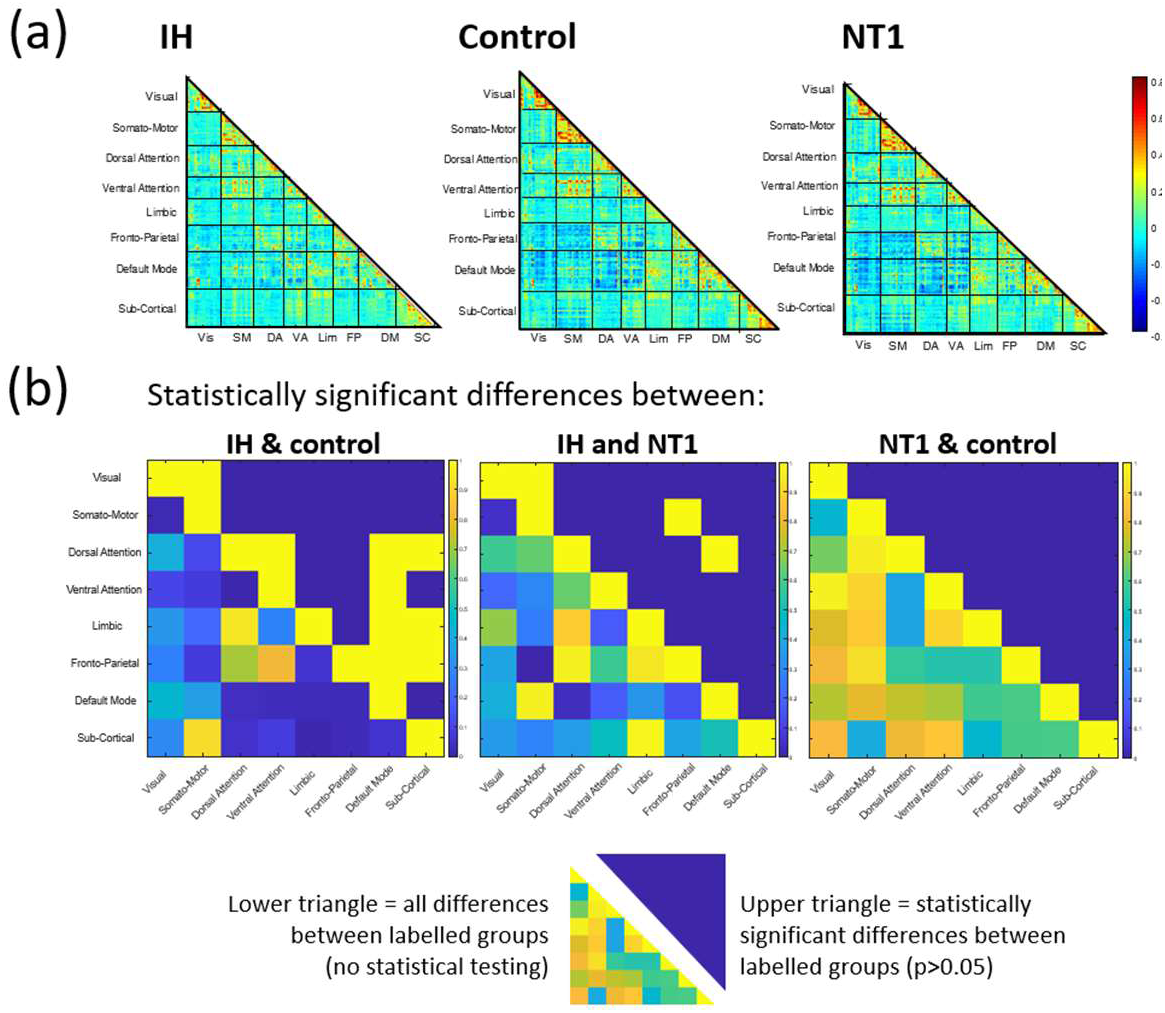
Summary figure of static results at baseline condition for all groups, at a network-level. (a) depicts total mean functional connectivity for all 246 ROIs across Yeo’s 7 networks and subcortical areas, warmer colors demonstrate higher subcortical connectivity in the NT1 group than the IH and control groups, and lower overall connectivity observed in the IH group compared to the other groups, (b) depicts the difference in mean functional connectivity across groups at baseline (lower triangle) and statistically significant differences (upper triangle). The patterns seen in (a) are reiterated here, though few significant (prior to multiple comparisons correction) differences across groups were found.

To complement the functional connectivity analyses, we then examined QPPs (Majeed et al. 2011). 2-D heatmap representations of the primary whole-brain pattern organized into Yeo’s 7 networks can be seen in Figure 3(a). There is clear anticorrelation occurring between network regions for all groups, though the individuals with NT1 have a noisier, less well-defined pattern than those with IH and the controls, as well as a much higher level of subcortical area activation during the QPP. The IH group QPP seems to have less overall activation at a network-level, as compared to the other groups. To interrogate this pattern further, the heatmap is translated onto a single-cycle waveform depiction in each group for networks of interest, DMN and TPN, shown in figure 3(b). In this representation, the TPN shows little activation (depicted by consistent low amplitude) in the control group, while the DMN oscillates more strongly and remains anticorrelated to it. This is measurably different than both the IH and NT1 group QPP, both of which have an overall higher level of TPN activation, and a dampened DMN amplitude. The control group’s low amplitude TPN is weakly correlated with the DMN, with a correlation coefficient of *0.20 (refer to Table 1),* while the IH group averages a DMN/TPN correlation of 0.33 – also positively correlated, though stronger than the control group value. The TPN is more active in the NT1 group, and we find a DMN/TPN correlation coefficient to be nearly uncorrelated, at −0.031. These correlation values are driven primarily by subnetworks, so those values (subnetwork correlations with DMN) are also included in *Table 1*, and plotted in figure 3(d). The squared difference in DMN/TPN per group was also plotted and is shown in figure 3(c), with the highest difference across conditions seen in the control group. This is not a statistical test but is included to further visualize the different functional organization observed across these groups. The correlation values plotted in figure 3(d) reveal the driving force behind the almost zero DMN/TPN correlation value observed from the NT1 group: their DAN/FPCNs are strongly correlated, compared to the other groups, which both demonstrate a moderately strong anticorrelation between these two networks. The IH group subnetwork correlation values are relatively close to the control group values, though are often weaker correlatively than the control.

**Figure 3:**
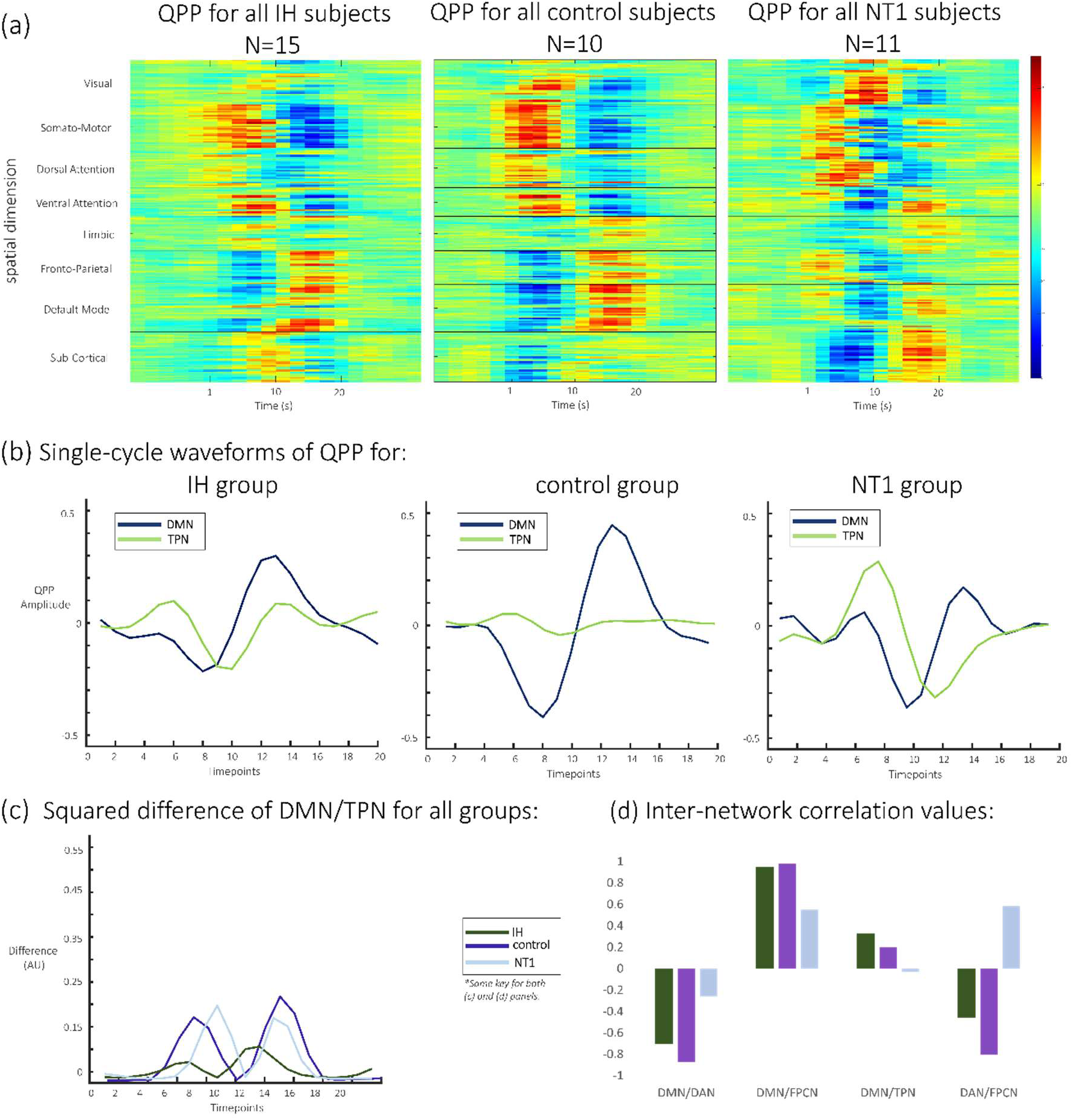
Summary figure of QPP results at baseline condition for all groups. (a) heatmap depictions of QPPs for each baseline group showing average activation for each of Yeo’s networks (labelled on y-axis) across the entire duration of the QPP (labelled on x-axis), demonstrating a clear visual difference between each group QPP, (b) waveform depiction of one cycle of the QPP for each group (DMN and TPN regions), where the control QPP is as expected, while the IH and NT1 groups show an altered, more correlated DMN/TPN QPP, (c) squared difference in TPN/DMN for all groups across one QPP cycle, showing the most difference in the control and NT1 groups, with little difference observed from the IH group, and (d) plotted inter-network correlation values for DMN and TPN composite networks (DAN, FPCN). These values are also included in table 1, and depict similar trends for both the IH and control groups, while the NT1 group differs in every comparison.

### Across Conditions Comparison

We then compared each group’s baseline condition to their results (functional connectivity and dynamic QPP) after a nap. For the comparisons of no-nap and post-nap conditions, 31 participants were included (IH n=12, NT1 n=11, control n=8), after excluding those participants who napped for less than 10 minutes and 1 participant who experienced a prolonged delay in transportation from the nap room to the MRI scanner. Figures 4 and 5 summarize the functional connectivity comparisons and analyses across conditions (upper triangle = nap condition, lower triangle = baseline condition) for all groups, finding ROI-level differences. Most notably, the NT1 and IH groups have overall weaker correlation between networks than the control group. Statistical significance was tested across groups and conditions at both an ROI-level (supplemental figure 4) and network-level (figure 5), though none survived multiple comparison correction. No group demonstrated significant difference across conditions, though there were some significant, uncorrected differences detected between groups.

**Figure 4:**
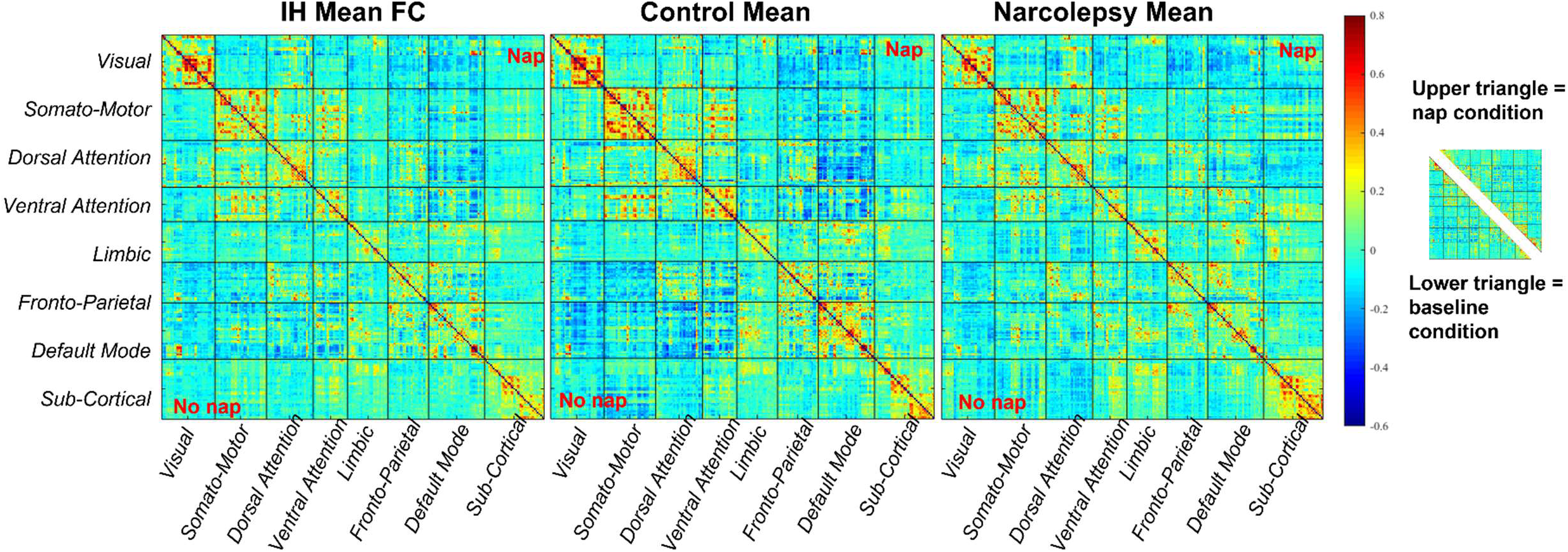
Average functional connectivity (scored via Pearson’s correlation coefficients) for all network regions for (L-R) IH, control and NT1 groups for both conditions (upper triangle = nap, lower triangle = baseline). While there are some differences to be observed across conditions here, it is difficult to identify which localized regions or networks are most affected.

**Figure 5:**
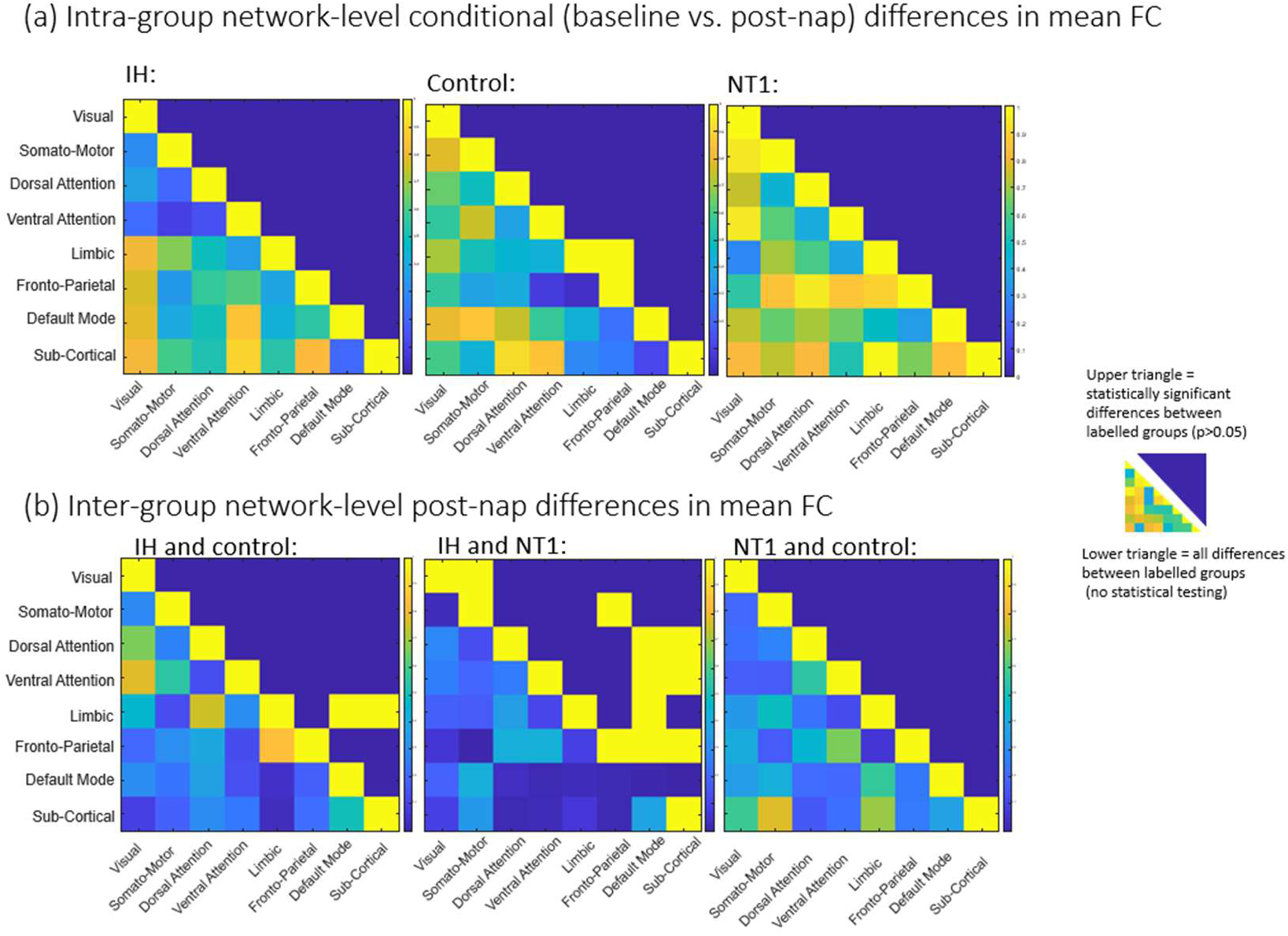
static results (mean functional connectivity) matrices for (a) comparison across conditions within groups and (b) comparison across groups for the post-nap differences in connectivity. For both (a) and (b), the upper triangle is the statistically significant values (not corrected for multiple comparisons; no values surpass that threshold), and the lower triangle is all differences across groups’ z-score per ROI. Almost no networks demonstrate a significant change in FC across conditions, though more are observed when comparing groups (in panel (b)).

QPP heatmaps for all 6 conditions (baseline and post-nap for the 3 groups) are depicted in Figure 6 for all 7 networks and subcortical regions. There are differences in network-level correlation both across groups and within a group across conditions. For example, the IH group at baseline has a less active DAN compared to the post-nap condition. However, at both conditions, the DMN and DAN are clearly anticorrelated – a trend also shown in the control for both conditions. This pattern is also somewhat evident in the NT1 group, but their QPP template shows a more disorganized pattern, with less clear anticorrelation between networks of interest. To quantify these differences, we then flattened the QPP to a single cycle over the window length (∼24s typically, 20s in this study) (Yousefi and Keilholz 2021), shown in figure 7.

**Figure 6:**
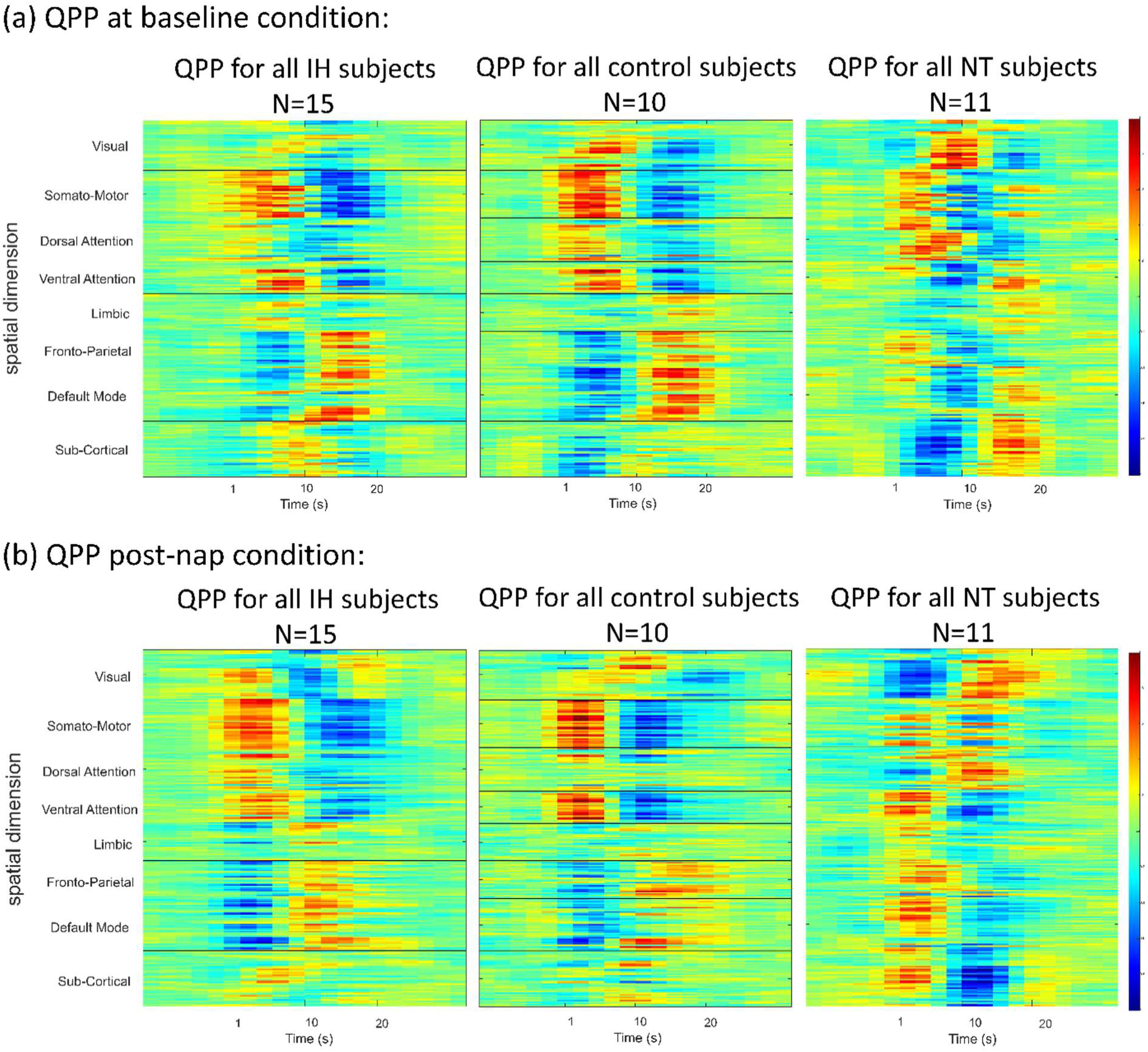
2-D representation of QPPs for each group and condition. (a) Each group QPP at baseline. (b) Each group QPP post-nap. All groups demonstrate some level of anticorrelation over time, particularly in Dorsal Attention network (DAN) regions and DMN regions, though this anticorrelation appears to be weaker in the ‘sleepy’ groups, particularly NT1.

**Figure 7:**
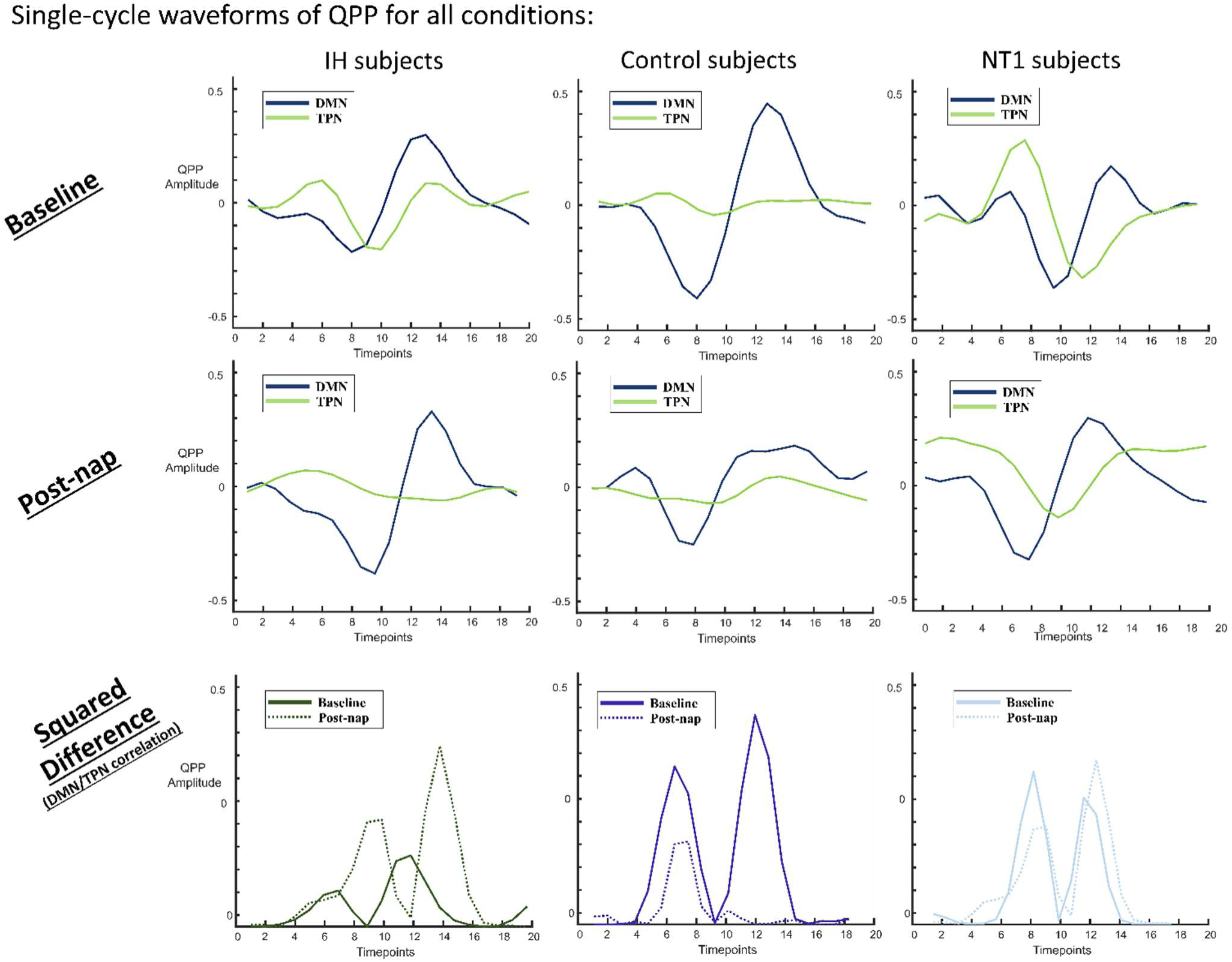
(a) Averaged QPP value per second for all groups (L-R: IH, control, NT1) at baseline conditions. The blue line (DMN) is typically strongly anticorrelated with the green line (TPN: DAN and FPCN) for the control group, less so for the IH and NT1 groups. The TPN in both the IH and NT1 groups is not plateaued like in the control group, but bimodal, implying TPN activation not seen in the control at baseline. *(b)* Averaged QPP value per second for all groups at nap condition, where all the groups show roughly the same DMN trend, though the TPN is vastly different for both the IH and NT1 groups. The TPN plateaus in the IH group, more closely mimicking the control group, while the NT1 group TPN shifts to a unimodal, strongly active TPN. *(c)* Squared difference of DMN and TPN for all groups (both conditions plotted on same axes), where only the NT1 group shows no change in the difference across conditions.

Looking at each group’s average QPP for only the DMN and TPN) is an effective way to visualize and quantify (amplitude, phase) the differences observed. Seen in figure 7, the IH group demonstrates a very consistent pattern in DMN activity across conditions – both in phase and amplitude range, a trend not seen in either of the groups. The TPN however does change, as it is significantly more active at rest than post-nap in the IH group, bringing the IH group closer to mimicking the control baseline waveform. The control group maintains an inactive TPN across conditions, but ultimately has a lower-amplitude DMN.

The NT1 group baseline QPP is notably dissimilar to the post-nap condition QPP, seeing a bimodal TPN at baseline and unimodal post-nap, though it is overall consistently active, particularly as compared to the control group. These DMN/TPN correlation values are included in Table 1, as well as other inter-network correlation values, with an * for statistical significance. The squared difference of the DMN/TPN for both conditions is plotted on a group-level, highlighting the range of variation that occurs within each group across conditions.

### Individual subject-level QPP detection

We observed some key changes across networks of interest, that are consistent with existing literature on vigilance signatures. No significance was observed within groups across conditions, perhaps due to the strong level of magnified noise and individual variability across the individual QPPs. However, there was significance detected when conducting a three-way comparison for each group across conditions. There was a statistically significant difference in DMN/TPN correlation post-nap, and nearly significant at baseline. These results are visualized in box plot form in figure 8. The three-way comparison of all groups also found significance at baseline for the DAN/FPCN correlation, though no significance is observed post-nap. These values and standard deviations are included in table 1 (italicized).

**Figure 8:**
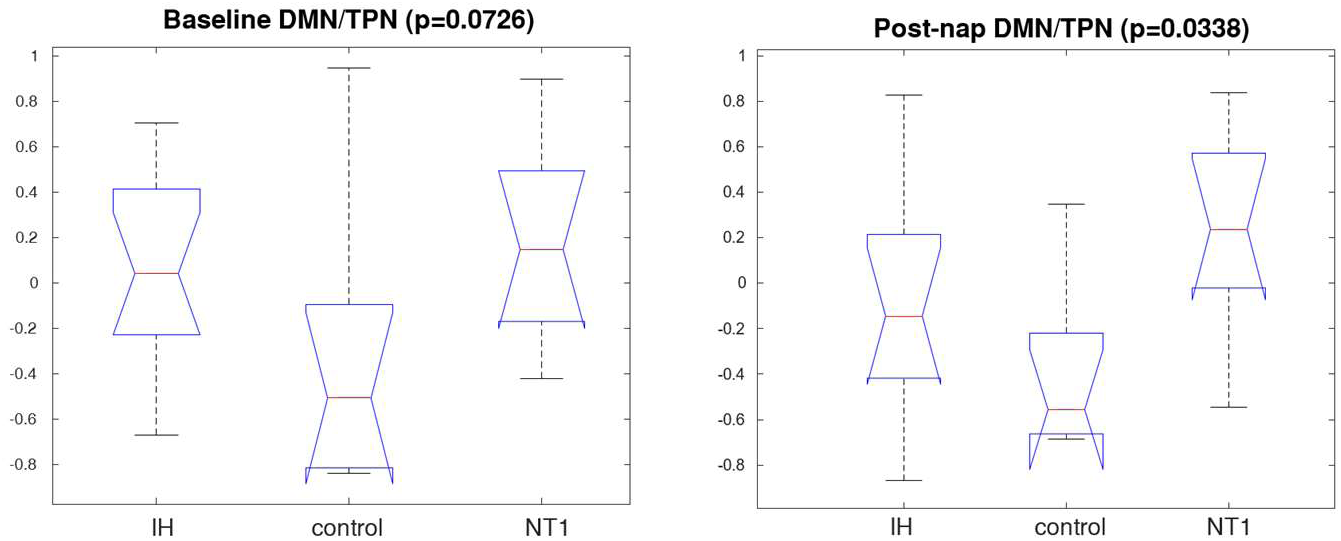
box plot of DMN/TPN correlation for all three groups’ individual QPPs, from a Kruskal-Wallis test (p>0.05). A significant difference across groups post-nap. The same analysis was conducted on other networks of interest (TPN subcomponents – DAN, FPCN) with respect to the DMN; these results are included in the supplemental figure 5. Other comparisons of individually calculated QPP inter-network correlation values are in supplemental figure 6.

### Global signal validation

The power spectra of global signal are plotted in figure 9. All groups have a higher global signal peak at baseline compared to post-nap, but the NT1 group consistently has the highest level at both conditions.

**Figure 9:**
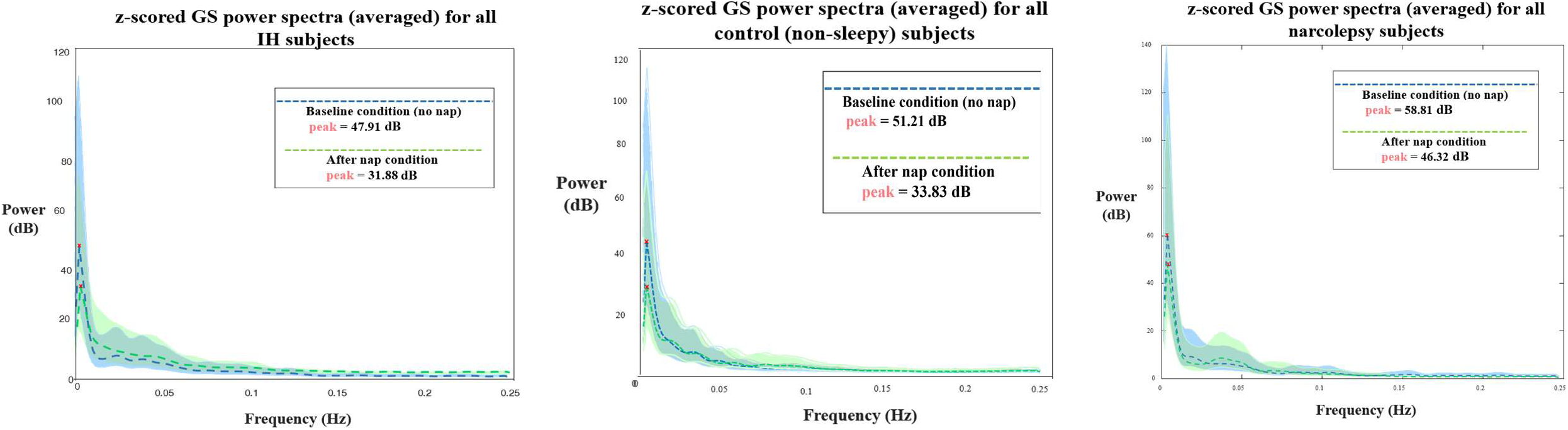
Z-scored global signal power spectra for both baseline (blue) and after nap (green) conditions for (L-R) IH, control and **NT1** subjects (all averaged). All peak in the infraslow band, but have varying levels of power at peak (values included on graphs). Dotted lines represent the group average, and the color-corresponding shading represents the group standard deviation.

### In-scanner sleep effects

We then considered the potential impacts of inadvertent sleep during the resting state scan on our results. For the baseline condition, simultaneous EEG data were available on 33 participants. Of these, at least one 30 second epoch of any stage of sleep occurred in 2 (15%) IH participants, 2 (18%) NT1 participants, and 1 (11%) control participant. When these data were removed, we continued to observe more overall connectivity in controls than in the patient groups and altered (increased) connectivity within subcortical regions in those with NT1 (Supplemental Figure 2(a)). There were no statistically significant differences in network-level connectivity between IH and the control group, but both IH and the control group had statistically significant somato-motor/FPCN connectivity when compared to the NT1 group – a finding consistent with other literature (Ruby et al. 2024). However, likely due to taking a small subset of an already initially small dataset, these differences are not significant after multiple comparison correction.From this subset of subjects, the dynamic analyses were able to reproduce many similar patterns observed from the entire dataset (supplemental figure 3). A few notable differences emerged, however, such as decreased DAN activity in the control group, decreased subcortical activity in the NT1 and IH groups, and affected TPN amplitude in all groups (supplemental figure 3(a)). As shown in the supplemental figure 3(b), the TPN amplitude weakened in both the IH and NT1 groups, but strengthened in the control group.

We planned to perform similar sensitivity analyses of sleep effects on the comparison between baseline and post-nap scans, by including those who were known to remain awake throughout both the baseline and post-nap scans. However, because sleep occurred in 6 participants in the post-nap scan (2 IH, 3 NT1, and 1 control), there was only a small subgroup of participants with no sleep on either scan (3 IH, 6 NT1, and 7 controls) and further subgroup analyses were not performed.

## Discussion

In this study, we demonstrated both distinct functional connectivity and alterations in QPPs in participants with IH and NT1 at baseline and post-nap, which can likely be attributed to the different disorder manifestations. The static functional connectivity results suggested target networks that were the *most* different across groups: the DMN, and TPN composite networks (DAN, FPCN). However, no statistically significant results were observed at a network-level for either condition, rendering static analysis challenging to use for analysis on its own. The dynamic analysis was employed to potentially identify any significance or notable patterns the static may have missed. We were able to pinpoint the primarily affected networks that drive the altered DMN/TPN correlation values seen in the IH and NT1 groups: while FPCN is more affected in the NT1 group, the DAN changes more across conditions in the IH group. Neither of these trends are reflected in the control group. Thus, the combination of the traditional static analysis, and the novel, dynamic QPP acquisition, provided a robust view of how functional organization is affected by IH and NT1 at baseline and after taking a nap.

As seen in figure 2(a), the overall mean FC is lower in both the IH and NT1 groups as compared to the controls, consistent with previously published studies, such as Pomares et al (2019). They conducted a comprehensive review of structural and functional differences in IH subjects compared to control subjects and found lower mean FC at rest between the anterior DMN and the orbitofrontal cortex network in people with IH compared to the non-sleepy control (defined as “good sleepers”) (Pomares et al. 2019). They also observed a negative correlation between mean FC and subjective daytime sleepiness. Fulong et al (2020) has also observed this relationship, in adolescents with NT1, in a published study which employed graph theory techniques to assess alterations in static FC (Fulong et al. 2020). They demonstrated decreased FC in the NT1 participants between the DMN regions and the left subcallosal gyrus, as well as increased FC in the visual network. There is little difference to report in the visual network FC of any of the three groups in this study, but there is notably more activity in sub-cortical regions in the group with NT1 compared to the other groups. This could be a result of instructing subjects to stay awake as long as possible - individuals with NT1 must try harder to stay awake, likely a process that is more demanding on sub-cortical structures (such as the hypothalamus) (Szymusiak, Gvilia, and McGinty 2007). This may also be why TPN regions in the NT1 group seems to be more active than the other groups, since they are potentially focusing more effort on staying awake which uses attentional networks. Other studies corroborate similar alterations in DMN FC and overall activation seen in individuals with NT1 (Gool et al. 2020; Engström et al. 2014; Järvelä et al. 2020). The IH group also shows a more activated TPN at baseline, which could also be explained as their trying harder to stay awake.

Comparing additional static measures of functional connectivity, both the IH group and the NT1 find no statistically significant differences at a network-level between conditions, and the only difference found from the control group is increased connectivity between the limbic network and the frontoparietal network, which does not survive multiple comparison correction. There are regions of significant difference found when comparing the IH group to the control, and NT1 for both conditions – at baseline, the IH group is statistically significantly different from the control group in the attentional networks. These networks are the DAN, VAN, FPCN, all of which have increased connectivity to the DMN; comparatively, post-nap, the only differences are within the limbic network connectivity to the DMN. None of these values survive multiple comparisons correction, but still provide useful information in identifying which networks are of interest in dynamic analysis. At baseline, there are several areas of significant difference between the IH group and the control group’s connectivity, decreased during the post-nap scan. This pattern is inverted when comparing the IH and NT1 groups, finding that ultimately individuals with both disorders respond to a nap differently. While there is some shared symptomatology between these two disorders – such as sleepiness and cognitive symptoms – the mechanistic differences may explain this variation of nap effect. Again, none of these comparative values survive multiple comparison analysis, emphasizing the need for complementary dynamic analysis to further explore and interrogate these data.

At resting-state, there is typically some level of DMN/TPN anticorrelation, not as strong as seen in task data due to an overall less active TPN, but still present (Abbas et al. 2019). This is seen in all groups, to varying degrees, particularly with the DMN and DAN, a key subnetwork of the TPN. As seen in supplemental figure 1, the DMN/DAN anticorrelation is likely strongest in the control group due to an overall stronger DMN amplitude, indicative of a “healthier” or more normal overall neural activity (Tang et al. 2017; Moran, Kelley, and Heatherton 2013). There is a much higher level of DAN activation seen in the NT1 group, as well as both the DAN and FPCN being out of phase. The FPCN is typically out of phase with the DMN during tasks and goal-oriented behavior, further supporting the claim that the subjects with NT1 were likely focusing their attention on something the other groups were not (such as trying not to fall asleep). The IH group also demonstrates a higher level of FPCN activation compared to the control across both conditions.

At baseline, people with IH and people with NT1 both suffer from severe sleepiness, so altered functional connectivity may in part reflect this shared experience of sleepiness. In contrast, after a nap, people with IH tend to still feel sleepy, while people with NT1 tend to feel more refreshed. Thus, a short nap provides a unique window for assessing for group differences in sleepiness across these two disorders. After a nap, the IH group’s QPP (as seen in figures 6-7) changes minimally, with a less active TPN, different from the drastic change seen in the NT1 group. This may reflect the restorative versus non-restorative effects of napping generally experienced by people with these two disorders. The minimal change in a single-cycle QPP DMN waveform, in conjunction with the self-reported feeling of minimal change before vs after a nap is unique to only those with IH, a key difference from the other groups and a key characteristic of the pathology. The less active TPN post-nap in the IH group does affect the DMN/TPN correlation value, yielding a moderate negative correlation between the two networks, somewhat similar to the NT1 baseline DMN/TPN correlation value. These patterns were mimicked when the QPP was detected at an individual subject level rather than a group aggregate; we expected to see high levels of variation and noise present, though we were still able to detect some significant differences (using p<0.05 as our threshold) – most notably the DMN/TPN correlation, which is nearly significant (p=0.073) at baseline, and significant (p=0.034) post-nap. This was further observed when we isolated the TPN subnetworks; the DAN particularly is starkly different in phase, amplitude and correlation to other networks across groups both at baseline and post-nap, observed with low p-values. The DAN has previously been used as a predictor of vigilance (Rosenberg et al. 2016), and is typically strongly anticorrelated with the DMN. Thus, the IH and NT1 groups likely have impaired DAN modulation, further validated by the DAN/FPCN significant finding at baseline.

The global signal power spectra (figure 9) was used to initially compare the IH and NT1 group results, as it is tightly inversely correlated to arousal level, so we expected both groups to differ from the control, as they are characterized by low arousal states. However, the IH group global signal was similar qualitatively and quantitatively to the control group at both conditions, and not uniquely change post-nap (compared to the control trend). The NT1 group global signal also followed the same baseline/post-nap trend of decreasing slightly, though the NT1 group demonstrated overall elevated signal across conditions, validating the low arousal state characterization.

To differentiate results between being disease-specific or simply a result of higher than average sleepiness, the subnetworks provide valuable insight. The sleepiness-dependent results are observed in similar trends in the NT1 and IH groups that are missing from the control group. Like the NT1 group, the IH group’s DMN/FPCN correlation was changed post-nap, whereas this value did not notably change across conditions in the control group. This change was not similar across groups however, as the NT1 group demonstrated a drastic increase in this correlation whereas the IH group showed a minor decrease. The FPCN is typically involved in executive function, goal-oriented tasks, and working memory tasks – all thought to be affected by poor sleep health (Yin, Li, and Chen 2022). If the IH group does not show as great of an effect on the FPCN post-nap as the NT1 group shows, this provides evidence to the symptom IH patients often report: that naps do not refresh them. Notable baseline differences across groups are seen in the DMN and DAN; both networks are less active in individuals with IH, a trait specific to the IH group. Since the DAN is typically responsible for externally directed tasks, it is possible that individuals with IH may focus more on introspection and internally directed tasks. It is also possible that those individuals’ brains may be modulating activity incorrectly, resulting in the excessive sleepiness, empowered by the underactive DAN during wakefulness.

There are still some confounds that need to be considered in parallel with the results and conclusions drawn. One that has been previously mentioned is that individuals with NT1 may have a harder time purposely staying awake in the MR scanner, particularly over the course of a 10-minute resting-state scan (as opposed to task-based, so not as engaging). Some participants in all three groups fell asleep in the scanner, despite specific instructions to try to remain awake, such that inadvertent sleep may account for some of the differences we see. However, our sensitivity analyses suggest this had a relatively modest effect on our results. There were also some technical limitations in the simultaneous EEG recordings, making EEG data unavailable for some participants. There was also some motion that occurred within included participants, which was not uniform across individuals, and as such, the homogenous motion correction applied may not have been sufficient for correcting this. It is also worth detailing that the nap conditions the subjects were under were non-ideal, as naps occurred near to but not within the MR scanner, and typical sleep lab-controlled conditions (e.g., light, noise) were not always available. Resting state scanning did not start immediately upon wakefulness, but after transportation from a room near to the scanner and completion of a 10-minute working memory task; some sleep inertia effects may have dissipated for some participants by the resting state scan.

## Conclusions

This study’s key findings can be used to better understand IH as a disease and how to potentially approach future mechanistic studies. This includes identifying baseline differences, such as the IH group having an underactive DAN on average, and conditional results, such as both sleepy groups demonstrating a differently correlated FPCN and DMN post-nap, not seen in the control group. This study also provides evidence for the anecdotal symptom IH patients often report through the unchanged QPP DMN amplitude: feeling unrefreshed after a nap, while typically healthy individuals report feeling better post-nap, as do individuals with NT1. The majority of these key findings stem from the dynamic analyses that preserve the temporal data, ensuring global *and* local patterns are detected. With only static analysis, the lack of significant findings may be discouraging, but used to identify target areas/networks for the dynamic analysis to focus on yielded a better understanding of how NT1 and IH differently affect an individual’s neural dynamic organization. These differences can be leveraged for investigating the root cause of this disease, and for future diagnostic purposes.

## Ethics

All data was acquired following Emory University Institutional Review Board.

## Data availability

Data and code will be made available upon request.

## Author contributions

LMT obtained funding for this study, and conceptualized and designed the study. PS and LMT acquired the data. HW and YB preprocessed and curated the data. LD performed the primary analyses and wrote the initial draft of the paper with LMT. SK and EH helped with analysis methodology and result interpretation. All authors helped with revision/editing of the manuscript.

## References

Abbas, Anzar, Yasmine Bassil, and Shella Keilholz. 2019. “Quasi-Periodic Patterns of Brain Activity in Individuals with Attention-Deficit/Hyperactivity Disorder.” NeuroImage : Clinical 21 (January):101653. 10.1016/j.nicl.2019.101653.

Abbas, Anzar, Michaël Belloy, Amrit Kashyap, Jacob Billings, Maysam Nezafati, Eric H. Schumacher, and Shella Keilholz. 2019. “Quasi-Periodic Patterns Contribute to Functional Connectivity in the Brain.” NeuroImage 191 (May):193–204. 10.1016/j.neuroimage.2019.01.076.

Allen, Elena A., Eswar Damaraju, Sergey M. Plis, Erik B. Erhardt, Tom Eichele, and Vince D. Calhoun. 2014. “Tracking Whole-Brain Connectivity Dynamics in the Resting State.” Cerebral Cortex (New York, NY) 24 (3): 663–76. 10.1093/cercor/bhs352.

Anumba, Nmachi, Eric Maltbie, Wen-Ju Pan, Theodore J. LaGrow, Nan Xu, and Shella Keilholz. 2023. “Spatial and Spectral Components of the BOLD Global Signal in Rat Resting-State Functional MRI.” Magnetic Resonance in Medicine 90 (6): 2486–99. 10.1002/mrm.29824.

Arand, Donna L., and Michael H. Bonnet. 2019. “The Multiple Sleep Latency Test.” Handbook of Clinical Neurology 160:393–403.

Arnulf, Isabelle, Smaranda Leu-Semenescu, and Pauline Dodet. 2022. “Precision Medicine for Idiopathic Hypersomnia.” Sleep Medicine Clinics 17 (3): 379–98.

Belloy, Michaël E., Maarten Naeyaert, Anzar Abbas, Disha Shah, Verdi Vanreusel, Johan van Audekerke, Shella D. Keilholz, Georgios A. Keliris, Annemie Van der Linden, and Marleen Verhoye. 2018. “Dynamic Resting State fMRI Analysis in Mice Reveals a Set of Quasi-Periodic Patterns and Illustrates Their Relationship with the Global Signal.” NeuroImage, Brain Connectivity Dynamics, 180 (October):463–84. 10.1016/j.neuroimage.2018.01.075.

Belloy, Michaël E., Disha Shah, Anzar Abbas, Amrit Kashyap, Steffen Roßner, Annemie Van der Linden, Shella D. Keilholz, Georgios A. Keliris, and Marleen Verhoye. 2018. “Quasi-Periodic Patterns of Neural Activity Improve Classification of Alzheimer’s Disease in Mice.” Scientific Reports 8 (1): 10024. 10.1038/s41598-018-28237-9.

Bola, Michał, and Viola Borchardt. 2016. “Cognitive Processing Involves Dynamic Reorganization of the Whole-Brain Network’s Functional Community Structure.” Journal of Neuroscience 36 (13): 3633–35.

Bolt, Taylor, Jason S. Nomi, Danilo Bzdok, Jorge A. Salas, Catie Chang, B. T. Thomas Yeo, Lucina Q. Uddin, and Shella D. Keilholz. 2022. “A Parsimonious Description of Global Functional Brain Organization in Three Spatiotemporal Patterns.” Nature Neuroscience 25 (8): 1093–1103. 10.1038/s41593-022-01118-1.

“BrainVision Analyzer | Brain Products GmbH > Solutions.” 2020. 2020. https://www.brainproducts.com/solutions/analyzer/.

Castellanos, F. Xavier, Daniel S. Margulies, Clare Kelly, Lucina Q. Uddin, Manely Ghaffari, Andrew Kirsch, David Shaw, Zarrar Shehzad, Adriana Di Martino, and Bharat Biswal. 2008. “Cingulate-Precuneus Interactions: A New Locus of Dysfunction in Adult Attention-Deficit/Hyperactivity Disorder.” Biological Psychiatry 63 (3): 332–37.

Craddock, Cameron, Sharad Sikka, Brian Cheung, Ranjeet Khanuja, Satrajit S. Ghosh, Chaogan Yan, Qingyang Li, Daniel Lurie, Joshua Vogelstein, and Randal Burns. 2013. “Towards Automated Analysis of Connectomes: The Configurable Pipeline for the Analysis of Connectomes (c-Pac).” Front Neuroinform 42 (10.3389). https://www.frontiersin.org/10.3389/conf.fninf.2013.09.00042/event_abstract?ref=https://githubhelp.com.

Dauvilliers, Yves, Elisa Evangelista, Lucie Barateau, Regis Lopez, Sofiène Chenini, Caroline Delbos, Séverine Beziat, and Isabelle Jaussent. 2019. “Measurement of Symptoms in Idiopathic Hypersomnia: The Idiopathic Hypersomnia Severity Scale.” Neurology 92 (15). 10.1212/WNL.0000000000007264.

Di, Xin, and Bharat B. Biswal. 2014. “Modulatory Interactions between the Default Mode Network and Task Positive Networks in Resting-State.” PeerJ 2:e367.

Diedenhofen, Birk, and Jochen Musch. 2015. “Cocor: A Comprehensive Solution for the Statistical Comparison of Correlations.” PloS One 10 (4): e0121945.

Du, Yuhui, Godfrey D. Pearlson, Qingbao Yu, Hao He, Dongdong Lin, Jing Sui, Lei Wu, and Vince D. Calhoun. 2016. “Interaction among Subsystems within Default Mode Network Diminished in Schizophrenia Patients: A Dynamic Connectivity Approach.” Schizophrenia Research 170 (1): 55–65.

Engström, Maria, Tove Hallböök, Attila Szakacs, Thomas Karlsson, and Anne-Marie Landtblom. 2014. “Functional Magnetic Resonance Imaging in Narcolepsy and the Kleine–Levin Syndrome.” Frontiers in Neurology 5:105.

Es, Mats WJ van, Cameron Higgins, Chetan Gohil, Andrew J. Quinn, Diego Vidaurre, and Mark W. Woolrich. 2023. “Large-Scale Cortical Networks Are Organized in Structured Cycles.” bioRxiv, 2023–07.

“False Discovery Rate.” 2016. Columbia University Mailman School of Public Health. August 8, 2016. https://www.publichealth.columbia.edu/research/population-health-methods/false-discovery-rate.

Fan, Lingzhong, Hai Li, Junjie Zhuo, Yu Zhang, Jiaojian Wang, Liangfu Chen, Zhengyi Yang, et al. 2016. “The Human Brainnetome Atlas: A New Brain Atlas Based on Connectional Architecture.” Cerebral Cortex 26 (8): 3508–26. 10.1093/cercor/bhw157.

Fulong, Xiao, Karen Spruyt, Lu Chao, Zhao Dianjiang, Zhang Jun, and Han Fang. 2020. “Resting-State Brain Network Topological Properties and the Correlation with Neuropsychological Assessment in Adolescent Narcolepsy.” Sleep 43 (8): zsaa018.

Gool, Jari K., Nathan Cross, Rolf Fronczek, Gert Jan Lammers, Ysbrand D. Van Der Werf, and Thien Thanh Dang-Vu. 2020. “Neuroimaging in Narcolepsy and Idiopathic Hypersomnia: From Neural Correlates to Clinical Practice.” Current Sleep Medicine Reports 6 (4): 251–66. 10.1007/s40675-020-00185-9.

Grooms, Joshua K., Garth J. Thompson, Wen-Ju Pan, Jacob Billings, Eric H. Schumacher, Charles M. Epstein, and Shella D. Keilholz. 2017. “Infraslow Electroencephalographic and Dynamic Resting State Network Activity.” Brain Connectivity 7 (5): 265–80. 10.1089/brain.2017.0492.

Huang, Yu-Shu, Ing-Tsung Hsiao, Feng-Yuan Liu, Fang-Ming Hwang, Kuang-Lin Lin, Wen-Cheng Huang, and Christian Guilleminault. 2018. “Neurocognition, Sleep, and PET Findings in Type 2 vs Type 1 Narcolepsy.” Neurology 90 (17): e1478–87. 10.1212/WNL.0000000000005346.

Järvelä, M., V. Raatikainen, A. Kotila, J. Kananen, V. Korhonen, L. Q. Uddin, H. Ansakorpi, and V. Kiviniemi. 2020. “Lag Analysis of Fast fMRI Reveals Delayed Information Flow between the Default Mode and Other Networks in Narcolepsy.” Cerebral Cortex Communications 1 (1): tgaa073.

Keilholz, Shella. 2023. “Time-Varying Functional Connectivity.” In Advances in Resting-State Functional MRI, 277–96. Elsevier. https://www.sciencedirect.com/science/article/pii/B9780323916882000060.

Kiviniemi, Vesa, Tapani Vire, Jukka Remes, Ahmed Abou Elseoud, Tuomo Starck, Osmo Tervonen, and Juha Nikkinen. 2011. “A Sliding Time-Window ICA Reveals Spatial Variability of the Default Mode Network in Time.” Brain Connectivity 1 (4): 339–47. 10.1089/brain.2011.0036.

Lee, Sangseok, and Dong Kyu Lee. 2018. “What Is the Proper Way to Apply the Multiple Comparison Test?” Korean Journal of Anesthesiology 71 (5): 353–60.

Liu, Xiao, Nanyin Zhang, Catie Chang, and Jeff H. Duyn. 2018. “Co-Activation Patterns in Resting-State fMRI Signals.” NeuroImage 180 (Pt B): 485–94. 10.1016/j.neuroimage.2018.01.041.

Lu, Jun, Thomas C. Jhou, and Clifford B. Saper. 2006. “Identification of Wake-Active Dopaminergic Neurons in the Ventral Periaqueductal Gray Matter.” Journal of Neuroscience 26 (1): 193–202.

Majeed, Waqas, Matthew Magnuson, Wendy Hasenkamp, Hillary Schwarb, Eric H. Schumacher, Lawrence Barsalou, and Shella D. Keilholz. 2011. “Spatiotemporal Dynamics of Low Frequency BOLD Fluctuations in Rats and Humans.” NeuroImage 54 (2): 1140–50. 10.1016/j.neuroimage.2010.08.030.

Maski, Kiran, Lynn Marie Trotti, Suresh Kotagal, R. Robert Auger, James A. Rowley, Sarah D. Hashmi, and Nathaniel F. Watson. 2021. “Treatment of Central Disorders of Hypersomnolence: An American Academy of Sleep Medicine Clinical Practice Guideline.” Journal of Clinical Sleep Medicine 17 (9): 1881–93. 10.5664/jcsm.9328.

Miglis, Mitchell G., Logan Schneider, Paul Kim, Joseph Cheung, and Lynn Marie Trotti. 2020. “Frequency and Severity of Autonomic Symptoms in Idiopathic Hypersomnia.” Journal of Clinical Sleep Medicine 16 (5): 749–56. 10.5664/jcsm.8344.

Moran, Joseph M., William M. Kelley, and Todd F. Heatherton. 2013. “What Can the Organization of the Brain’s Default Mode Network Tell Us about Self-Knowledge?” Frontiers in Human Neuroscience 7:391.

Pomares, Florence B., Soufiane Boucetta, Francis Lachapelle, Jason Steffener, Jacques Montplaisir, Jungho Cha, Hosung Kim, and Thien Thanh Dang-Vu. 2019. “Beyond Sleepy: Structural and Functional Changes of the Default-Mode Network in Idiopathic Hypersomnia.” Sleep 42 (11): zsz156.

Raichle, Marcus E., Ann Mary MacLeod, Abraham Z. Snyder, William J. Powers, Debra A. Gusnard, and Gordon L. Shulman. 2001. “A Default Mode of Brain Function.” Proceedings of the National Academy of Sciences 98 (2): 676–82. 10.1073/pnas.98.2.676.

Ramar, Kannan, Raman K. Malhotra, Kelly A. Carden, Jennifer L. Martin, Fariha Abbasi-Feinberg, R. Nisha Aurora, Vishesh K. Kapur, et al. 2021. “Sleep Is Essential to Health: An American Academy of Sleep Medicine Position Statement.” Journal of Clinical Sleep Medicine 17 (10): 2115–19. 10.5664/jcsm.9476.

Ramm, Markus, Matthias Boentert, Nelly Lojewsky, Arsalan Jafarpour, Peter Young, and Anna Heidbreder. 2019. “Disease-Specific Attention Impairment in Disorders of Chronic Excessive Daytime Sleepiness.” Sleep Medicine 53:133–40.

Rassu, Anna Laura, Elisa Evangelista, Lucie Barateau, Sofiene Chenini, Régis Lopez, Isabelle Jaussent, and Yves Dauvilliers. 2022. “Idiopathic Hypersomnia Severity Scale to Better Quantify Symptoms Severity and Their Consequences in Idiopathic Hypersomnia.” Journal of Clinical Sleep Medicine 18 (2): 617–29. 10.5664/jcsm.9682.

Rosenberg, Monica D., Emily S. Finn, Dustin Scheinost, Xenophon Papademetris, Xilin Shen, R. Todd Constable, and Marvin M. Chun. 2016. “A Neuromarker of Sustained Attention from Whole-Brain Functional Connectivity.” Nature Neuroscience 19 (1): 165–71. 10.1038/nn.4179.

Ruby, Perrine, Elisa Evangelista, Hélène Bastuji, and Laure Peter-Derex. 2024. “From Physiological Awakening to Pathological Sleep Inertia: Neurophysiological and Behavioural Characteristics of the Sleep-to-Wake Transition.” Neurophysiologie Clinique 54 (2): 102934.

Salvador, Raymond, Erick Canales-Rodríguez, Amalia Guerrero-Pedraza, Salvador Sarró, Diana Tordesillas- Gutiérrez, Teresa Maristany, Benedicto Crespo-Facorro, Peter McKenna, and Edith Pomarol-Clotet. 2019. “Multimodal Integration of Brain Images for MRI-Based Diagnosis in Schizophrenia.” Frontiers in Neuroscience 13:1203.

Sateia, Michael J. 2014. “International Classification of Sleep Disorders-Third Edition: Highlights and Modifications.” Chest 146 (5): 1387–94. 10.1378/chest.14-0970.

Satpute, Ajay B., and Kristen A. Lindquist. 2019. “The Default Mode Network’s Role in Discrete Emotion.” Trends in Cognitive Sciences 23 (10): 851–64.

Specht, Karsten. 2020. “Current Challenges in Translational and Clinical fMRI and Future Directions.” Frontiers in Psychiatry 10:924.

Szymusiak, Ronald, Irma Gvilia, and Dennis McGinty. 2007. “Hypothalamic Control of Sleep.” Sleep Medicine 8 (4): 291–301.

Takahashi, Masaya. 2003. “The Role of Prescribed Napping in Sleep Medicine.” Sleep Medicine Reviews 7 (3): 227–35.

Tang, Wei, Hesheng Liu, Linda Douw, Mark A. Kramer, Uri T. Eden, Matti S. Hämäläinen, and Steven M. Stufflebeam. 2017. “Dynamic Connectivity Modulates Local Activity in the Core Regions of the Default-Mode Network.” Proceedings of the National Academy of Sciences 114 (36): 9713–18. 10.1073/pnas.1702027114.

Thompson, Garth John, Wen-Ju Pan, Matthew Evan Magnuson, Dieter Jaeger, and Shella Dawn Keilholz. 2014. “Quasi-Periodic Patterns (QPP): Large-Scale Dynamics in Resting State fMRI That Correlate with Local Infraslow Electrical Activity.” NeuroImage 84 (January):1018–31. 10.1016/j.neuroimage.2013.09.029.

Trotti, Lynn Marie. 2017. “Idiopathic Hypersomnia.” Sleep Medicine Clinics 12 (3): 331–44.

Trotti, Lynn Marie, Prabhjyot Saini, Donald L. Bliwise, Amanda A. Freeman, Andrew Jenkins, and David B. Rye. 2015. “Clarithromycin in Γ-aminobutyric Acid–Related Hypersomnolence: A Randomized, Crossover Trial.” Annals of Neurology 78 (3): 454–65. 10.1002/ana.24459.

Uddin, Lucina Q., AM Clare Kelly, Bharat B. Biswal, Daniel S. Margulies, Zarrar Shehzad, David Shaw, Manely Ghaffari, John Rotrosen, Lenard A. Adler, and F. Xavier Castellanos. 2008. “Network Homogeneity Reveals Decreased Integrity of Default-Mode Network in ADHD.” Journal of Neuroscience Methods 169 (1): 249–54.

Wang, Kai, Waqas Majeed, G. J. Thompson, Kui Ying, Yan Zhu, and S. D. Keilholz. 2016. “Quasi-Periodic Pattern of fMRI Contributes to Functional Connectivity and Explores Differences between Major Depressive Disorder and Control.” In Proc Int Soc Magn Reson Med. Vol. 1683. https://archive.ismrm.org/2016/1683.html.

Wu, Lanxiang, Qingqing Zhan, Qian Liu, Suheng Xie, Sheng Tian, Liang Xie, and Wei Wu. 2022. “Abnormal Regional Spontaneous Neural Activity and Functional Connectivity in Unmedicated Patients with Narcolepsy Type 1: A Resting-State Fmri Study.” International Journal of Environmental Research and Public Health 19 (23): 15482.

Yin, Shouhang, Yilu Li, and Antao Chen. 2022. “Functional Coupling between Frontoparietal Control Subnetworks Bridges the Default and Dorsal Attention Networks.” Brain Structure and Function 227 (7): 2243–60. 10.1007/s00429-022-02517-7.

Yousefi, Behnaz, and Shella Keilholz. 2021. “Propagating Patterns of Intrinsic Activity along Macroscale Gradients Coordinate Functional Connections across the Whole Brain.” NeuroImage 231 (May):117827. 10.1016/j.neuroimage.2021.117827.

