## Supplemental Information for "Altered functional connectivity and spatiotemporal dynamics in individuals with sleep disorders"

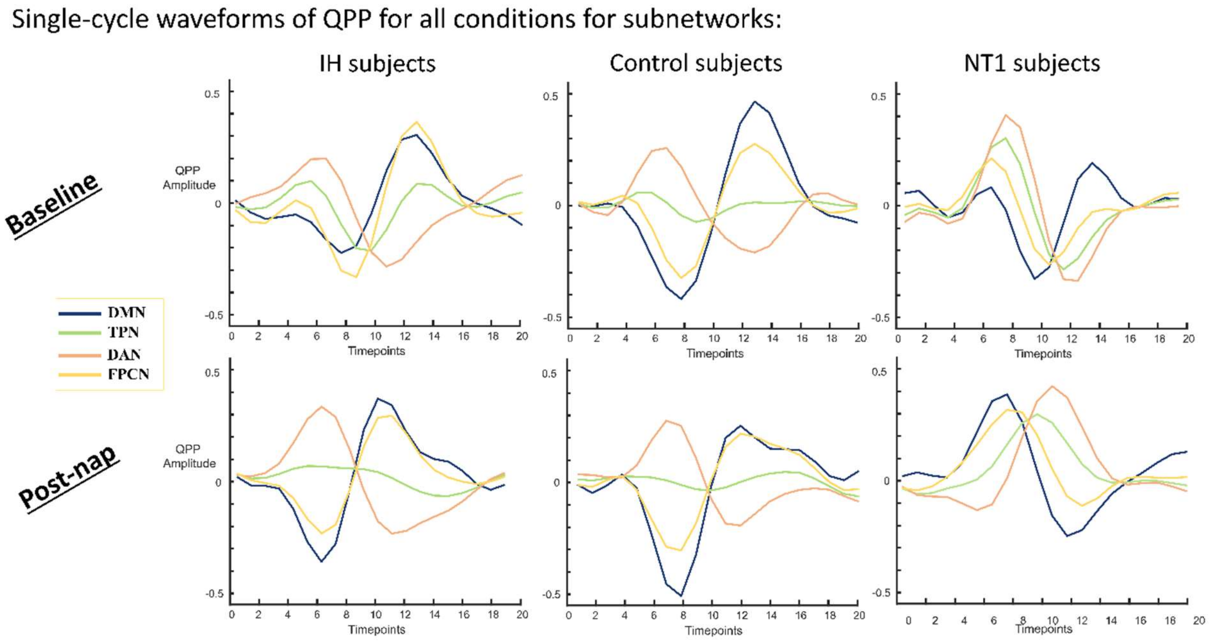


Supplemental figure 1 (S1): single-cycle waveforms of QPP for DMN, TPN and TPN composite networks (DAN, FPCN).

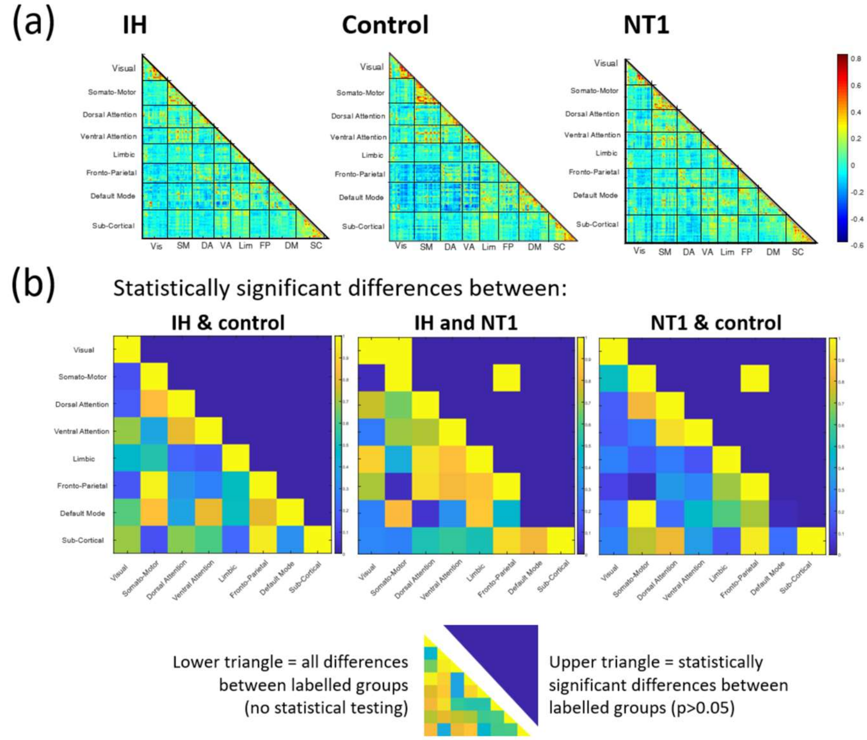


Supplemental figure 2 (S2): static results from all groups' baseline condition for a subset of subjects (those who definitely did not fall asleep in the scanner: healthy N=8, IH N=10, NT1 N=7), (a)

connectivity matrices for all three groups, (b) statistical significance results, not controlled for multiple comparisons (no values survived so the uncontrolled values are reported here).

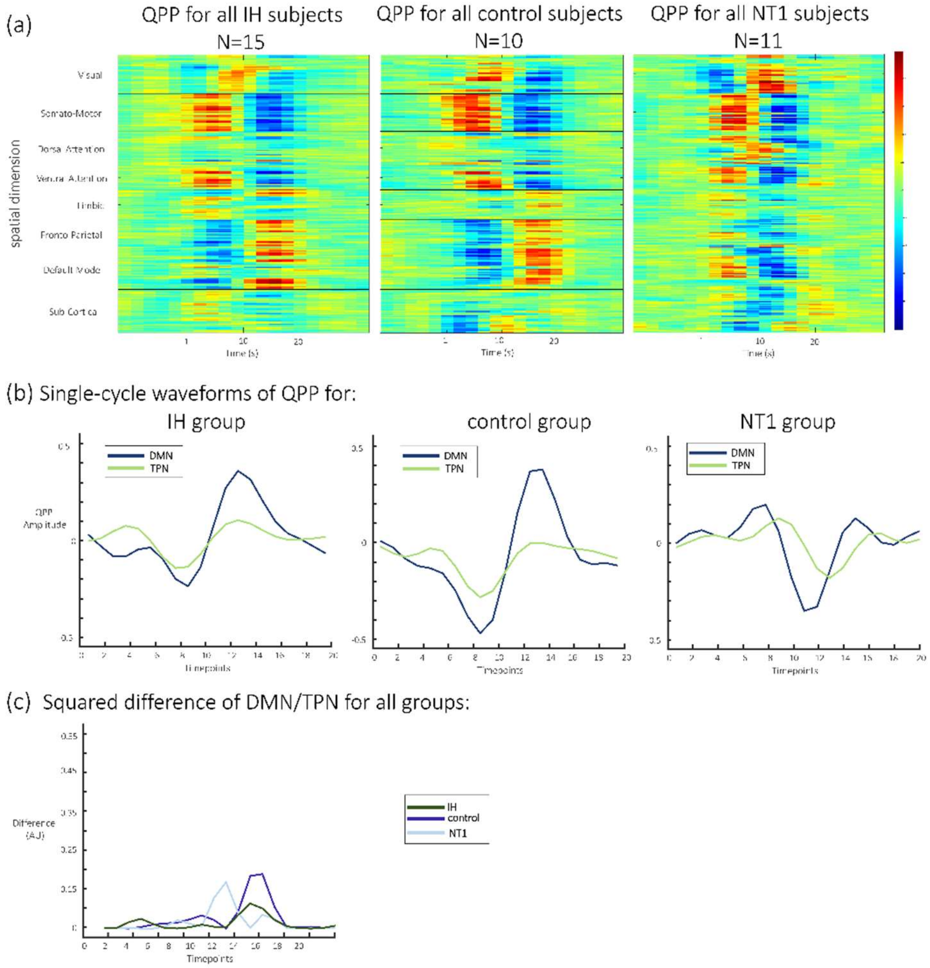


Supplemental figure 3 (S3): dynamic results from all groups' baseline condition for a subset of subjects (those who definitely did not fall asleep in the scanner), (a) QPP heatmaps, (b) single-cycle waveforms of QPP, (c) squared difference of DMN/TPN correlation for all groups on the same axis

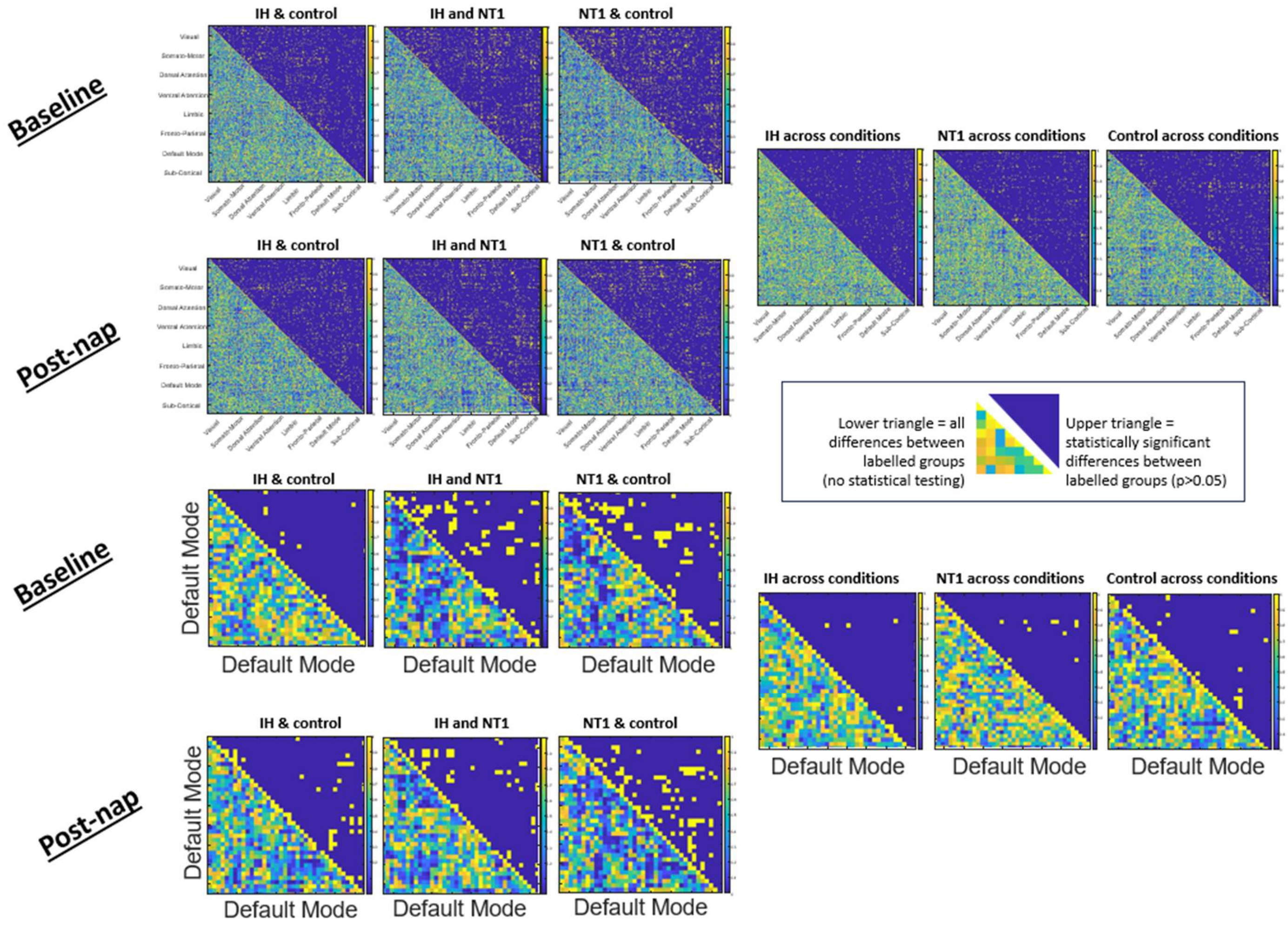


Supplemental figure 4 (S4): static connectivity results for all conditions at an ROI-level, for both all networks and the DMN ROIs. All rows are labeled for which groups/conditions are compared, the legend (upper/lower triangle) applies for all.

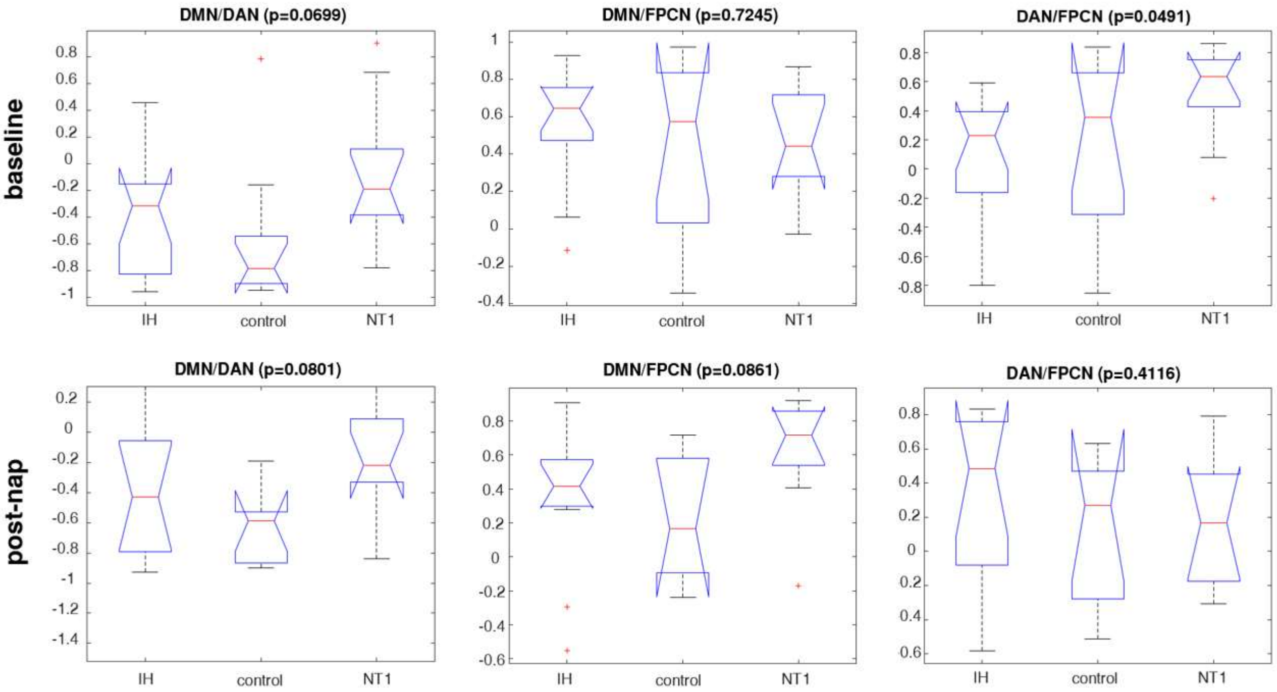


Supplemental figure 5 (S5): From the individually observed QPPs, these are box plots of inter-network correlations for all three groups, from a Kruskal-wallis test (p>0.05 for significance). The first row is DMN/DAN, DMN/FPCN and DAN/FPCN correlation values at baseline condition; the second row is the same correlation value groups at the post-nap condition.

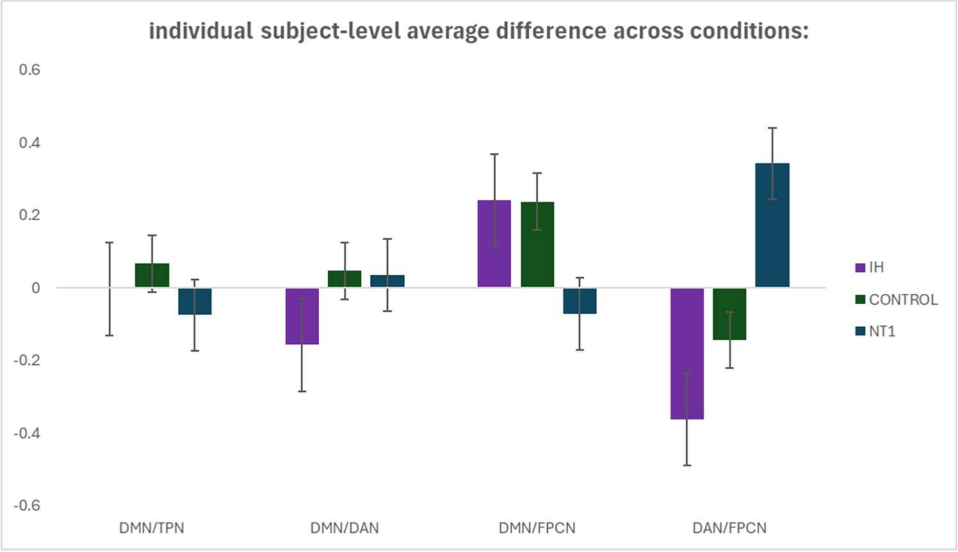


Supplemental figure 6 (S6): From the individually observed QPPs, this plot compares the inter-network correlation values for all groups, with error bars.
